# Pre-pregnant obesity of mothers in a multi-ethnic cohort is associated with cord blood metabolomic changes in offspring

**DOI:** 10.1101/264374

**Authors:** Ryan J. Schlueter, Fadhl M. Al-Akwaa, Paula A. Benny, Alexandra Gurary, Guoxiang Xie, Wei Jia, Shaw J. Chun, Ingrid Chern, Lana X. Garmire

## Abstract

Maternal obesity has become a growing global health concern that may predispose the offspring to medical conditions later in life. However, the metabolic link between maternal pre-pregnant obesity and healthy offspring has not yet been fully elucidated. In this study, we conducted a case-control study using coupled untargeted and targeted metabolomics approach, from the newborn cord blood metabolomes associated with a matched maternal pre-pregnant obesity cohort of 28 cases and 29 controls. The subjects were recruited from multi-ethnic populations in Hawaii, including rarely reported Native Hawaiian and other Pacific Islanders (NHPI). We found that maternal obesity was the most important factor contributing to differences in cord blood metabolomics. Using elastic net regularization based logistic regression model, we identified 29 metabolites as potential early-life biomarkers manifesting intrauterine effect of maternal obesity, with accuracy as high as 0.947 after adjusting for clinical confounding (maternal and paternal age and ethnicity, parity and gravidity). We validated the model results in a subsequent set of samples (N=30) with an accuracy of 0.822. Among the metabolites, six metabolites (galactonic acid, butenylcarnitine, 2-hydroxy-3-methylbutyric acid, phosphatidylcholine diacyl C40:3, 1,5-anhydrosorbitol, and phosphatidylcholine acyl-alkyl 40:3) were individually and significantly different between the maternal obese vs. norm-weight groups. Interestingly, Hydroxy-3-methylbutyric acid showed significnatly higher levels in cord blood from the NHPI group, compared to asian and caucasian groups. In summary, significant associations were observed between maternal pre-pregnant obesity and offspring metabolomics alternation at birth, revealing the inter-generational impact of maternal obesity.

## Introduction

Obesity is a global health concern. While some countries have a relative paucity of obesity, in the United States, obesity affects 38% of adults ^1, 2^. As such, maternal obesity has risen to epidemic proportions in recent years and can impose significant risk to both the mother and unborn fetus. By 2015, an estimated 25.6 % women were obese before becoming pregnant, according to a CDC study^3^. Maternal pre-pregnant obesity can increase the risk for a wide range of health concerns for the baby and the mother during all stages of pregnancy. Moreover, research has recently extended the association of maternal obesity during pregnancy to the subsequent health of offspring such as diabetes or cardiovascular disease ^4^. Since the inception of Barker’s hypothesis in the 1990’s, efforts to connect intrauterine exposures with the development of disease later in life has been the subject of many studies ^5^. Both obesity and its accompanying morbidities, such as diabetes, cardiovascular diseases and cancers, are of particular interest as considerable evidence has shown that maternal metabolic irregularities may have a role in genotypic programming in offspring ^6, 7^. Identifying markers of predisposition to health concerns or diseases would present an opportunity for early identification and potential intervention, thus providing life-long benefits^8–10^.

Previous studies have found that infants born to obese mothers consistently demonstrate elevation of adiposity and are at more substantial risk for the development of metabolic disease ^11^. While animal models have been used to demonstrate early molecular programming under the effect of obesity, human research to elucidate the underlying mechanisms in origins of childhood disease is lacking ^12^. In *Drosophila melanogaster*, offspring of females given a high-sucrose diet exhibited metabolic aberrations both at the larvae and adult developmental stages ^13, 14^. Though an invertebrate model, mammalian lipid and carbohydrate systems show high level of conservation in *Drosophila melanogaster* ^15, 16^. In a mouse model of maternal obesity, progeny demonstrated significant elevations of both leptin and triglycerides when compared with offspring of control mothers of normal weight ^6^. The authors proposed that epigenetic modifications of obesogenic genes during intrauterine fetal growth play a role in adaption to an expected future environment. Recently, Tillery *et al.* used a primate model to examine the origins of metabolic disturbances and altered gene expression in offspring subjected to maternal obesity ^17^. The offspring consistently displayed significant increases in triglyceride level and also fatty liver disease on histologic preparations. However, human studies that explore the fetal metabolic consequences of maternal obesity are still in need of investigation.

Metabolomics is the study of small molecules using high throughput platforms, such as mass spectroscopy^18^. It is a desirable technology that can detect distinct chemical imprints in tissues and body fluids ^19^. The field of metabolomics has shown great promise in various applications including early diagnostic marker identification ^20^, where a set of metabolomics biomarkers can differentiate samples of two different states (eg. disease and normal states). Cord blood metabolites provide information on fetal nutritional and metabolic health ^21^, and could provide an early window of detection to potential health issues among newborns. Previously, some studies have reported differential metabolite profiles associated with pregnancy outcomes such as intrauterine growth restriction ^22^ and low birth weight ^23^. For example, abnormal lipid metabolism and significant differences in relative amounts of amino acids were found in metabolomic signatures in cord blood from infants with intrauterine growth restriction in comparison to normal weight infants ^22^. In another study higher phenylalanine and citrulline levels but lower glutamine, choline, alanine, proline and glucose levels were observed in cord blood of infants of low-birth weight ^23^. However, thus far no metabolomics studies have been reported to specifically investigate the impact of maternal obesity on metabolomics profiles in fetal cord blood ^22–25^.

This study aims to investigate metabolomics changes in fetal cord blood associated with obese (BMI>30) and normal pre-pregnant weight (18.5<BMI<25) mothers. Uniquely, we recruited mothers from the multi-ethnic population in Hawaii, including Native Hawaiian and other Pacific Islanders (NHPI). NHPI is a particularly under-represented minority population across most scientific studies.

## Methods

### Chemicals & Reagents

Ethanol, pyridine, methoxyamine hydrochloride, C8–C30 fatty acid methyl esters (FAMEs), and ammonium acetate were purchased from Sigma Aldrich (St. Louis, MO). LC-MS Optima grade methanol and acetonitrile, formic acid, N-Methyl-N-trimethylsilyltrifluoroacetamide (MSTFA) with 1% TMCS, and hexane were obtained from Fisher Sci. (Fair Lawn, NJ). The ultra-pure water was produced by Millipore Advantage A10 system with a LC-MS Polisher filter (Billerica, MA). Analytical grade sodium hydroxide, sodium bicarbonate, and anhydrous sodium sulfate were obtained from JT Baker Co. (Phillipsburg, NJ, USA). The amino acid and lipid standards were included in the AbsoluteIDQ™ p180 Kit (Biocrates Life Sciences, Austria). All other standards were commercially purchased from Sigma-Aldrich and Nu-Chek Prep (Elysian, MN, USA).

### Study population

We performed a multi-ethnic case-control study at Kapiolani Medical Center for Women and Children, Honolulu, HI from June 2015 through June 2017. The study was approved by the Western IRB board (WIRB Protocol 20151223). To avoid confounding of inflammation accompanying labor and natural births ^26^ we recruited women scheduled for full-term cesarean section at ≥ 37 weeks gestation. All subjects fasted for at least 8 hours before the scheduled cesarean delivery. Patients meeting inclusion criteria were identified from pre-admission medical records with pre-pregnancy BMI ≥30.0 (cases) or 18.5-25.0 (controls). The pre-pregnancy BMIs were also confirmed during the enrollment. Women with preterm rupture of membranes (PROM), labor (being active contractions with cervical dilation), multiple gestations, pre-gestational diabetes, hypertensive disorders, cigarette smokers, HIV, HBV, and chronic drug users were excluded. Clinical characteristics were recorded, including maternal and paternal age, maternal and paternal ethinicities, mother’s pre-pregnancy BMI, net weight gain, gestational age, parity, gravidity and ethnicity. Neonate weight was recorded in kilograms and the weight of baby was taken directly after the birth in the newborn nursey. For the discovery cohort, a total of 57 subjects (28 cases and 29 controls) were recruited. Additionally, to confirm the results, we recruited 30 subjects (12 cases and 18 controls) from the same site but different time interval (July 2017 to June 2018).

### Sample collection, preparation and quality control

Cord blood was collected under sterile conditions at the time of cesarean section using Pall Medical cord blood collection kit with 25 mL citrate phosphate dextrose (CPD) in the operating room. The umbilical cord was cleansed with chlorhexidine swab before collection to ensure sterility. The volume of collected blood was measured and recorded before aliquoting to conicals for centrifugation. Conicals were centrifuged at 200g for 10 minutes and plasma was collected. The plasma was centrifuged at 350g for 10 minutes and aliquoted into polypropylene cryotubes, and stored at −80C.

The investigators all took and passed courses where transport, collection and laboratory use of biologic specimens was tested. During the handling of samples gloves were used and documentation for biohazard materials accompanied the transportation of materials. Laboratory grade cryo-plasticware was used for storage and all samples were labeled stringently and kept in 100-well boxes with record sheets in Excel spreadsheet. Upon the use of samples for metabolome profiling, all samples were treated as biohazards. The investigators all received appropriate vaccinations and personal protective equipment including gloves, lab coats, disposable face mask, and glasses for eye protection were used during sample preparation.

### Metabolome profiling

The plasma samples were thawed and extracted with 3-vol cold organic mixture of ethanol: chloroform and centrifuged at 4 °C at 14,500 rpm for 20 min. The supernatant was split for lipid and amino acid profiling with an Acquity ultra performance liquid chromatography coupled to a Xevo TQ-S mass spectrometry (UPLC-MS/MS, Waters Corp., Milford, MA). Metabolic profiling of other metabolites including organic acids, carbohydrates, amino acids, and nucleotides were measured with an Agilent 7890A gas chromatography coupled to a Leco Pegasus time of flight mass spectrometry (Leco Corp., St Joseph, MI). The raw data files generated from LC-MS (targeted) and GC-MS (untargeted) were processed with TargetLynx Application Manager (Waters Corp., Milford, MA) and ChromaTOF software (Leco Corp., St Joseph, MI) respectively. Peak signal, mass spectral data, and retention times were obtained for each metabolite. The detected metabolites from GC-MS were annotated and combined using an automated mass spectral data processing software package ^27^. The levels of lipids and amino acids detected from LC-MS were measured using the AbsoluteIDQ™ p180 Kit (Biocrates Life Sciences, Austria) commercially available. The reference standards of those measured lipids and amino acids were integrated in the kit^28^. More details of metabolomics experiments and data pre-processing is described in the following subsections.

### Sample Preparation for metabolic profiling

Plasma samples were prepared as previously described with minor modifications^29^. Each 150 µL of cold organic mixture (ethanol: chloroform = 3:1, v/v) is used to extract small-molecule metabolites from 50 µL of blood sample, spiked with two internal standard solutions (10 µL of L-2-chlorophenylalanine in water, 0.3 mg/ml; 10 µL of heptadecanoic acid in methanol, 1 mg/mL). The sample extracts were centrifuged at 4 °C and 14, 500 rpm for 20 min. The supernatant was split for lipid and amino acid profiling with an Acquity ultra performance liquid chromatography coupled to a Xevo TQ-S mass spectrometry (UPLC-MS/MS, Waters Corp., Milford, MA) and for untargeted metabolic profiling with Gas Chromatography -Time-of-Flight Mass Spectrometry (GC−TOFMS).

### Quantitation of amino acids and lipids with LC-MS/MS

For targeted metabolomics analyses of plasma samples, the AbsoluteIDQ™ p180 Kit (Biocrates Life Sciences, Austria) was used, which allows for the simultaneous quantification of metabolites from different compound classes [21 amino acids (AA), 21 biogenic amines (BA), 40 acylcarnitines (AC), 38 acyl/acyl phosphatidylcholines (PC aa), 38 acyl/alkyl phosphatidylcholines (PC ae), 14 lyso-phosphatidylcholines (lysoPC), 15 sphingomyelins (SM), and the sum of hexoses (H1)]. The lipids, acylcarnitines, and the hexoses were determined by FIA-MS/MS, while the amino acids and biogenic amines were measured by LC-MS/MS. In brief, each aliquot of the 20-µL supernatant was added to a 96-well Biocrates Kit plate (Biocrates Life Sciences, Austria) for metabolite quantitation. After samples were dried under nitrogen, each 300 µL of extraction solvent (5 mM ammonium acetate in methanol) was added and the kit plate was gently shaken at room temperature for 30 min. The extracts were derivatizaed with phenylisothiocyanate for amino acids and biogenic amine quantification. The data was acquired by using MassLynx 4.1 software (Waters) and was analyzed using TargetLynx applications manager version 4.1 (Waters) to obtain calibration equations and the quantitative concentration of each metabolite in the samples. Another aliquot of 20 µL of the extracts was further diluted with 380 µL of methanol with 5 mM ammonium acetate for FIA analysis of lipids. An Acquity ultra performance liquid chromatography coupled to a Xevo TQ-S mass spectrometer (UPLC-MS/MS, Waters Corp., Milford, MA) was used for targeted metabolite analysis of 140 lipids in cell line samples. Each 10 µL of sample was directly injected into mass spectrometer with elution solvent (methanol with 5 mM ammonium acetate) at a varied flow rate from 30 to 200 µL/min within 3 min. ^30^ Concentrations of lipids were directly calculated in MetIDQ (version 4.7.2, Biocrates).

### Untargeted Metabolomics Profiling with GC-TOFMS

The protocol for untargeted metabolomics profiling was reported earlier ^31, 32^. Each aliquot of the 150-µL above supernatant was dried in a vacuum centrifuge concentrator. The dried material is derivatized with 50 µL of methoxyamine (15 mg/mL in pyridine) at 30°C for 90 min. After the addition of 10 μL of alkynes (retention index standards) and 50 μL of BSTFA (1% TMCS), the mixture is incubated at 70°C for 60 min. Retention indices of C8−C30 fatty acid methyl esters (FAMEs) were added for retention-time correction. Each 1-µl sample was analyzed on an Agilent 7890A gas chromatograph coupled to a Leco Pegasus time of flight mass spectrometer (Leco Corp., St Joseph, MI) for global metabolite analysis. The analytes were introduced with a splitless mode to achieve maximum sensitivity and separated on a Rtx-5 MS capillary column (30 m x 0.25 mm I.D., 0.25 µm) (Restek, Bellefonte, PA). The column temperature was initially set to 80 °C for 2 min, increased to 300 °C in 12 min, and maintained at 300 °C for 5 min.

The solvent delay was set to 4.4 min. The front inlet temperature, transferline temperature, and source temperature were set to 260 °C, 270 °C, and 220 °C, respectively. The mass spectrometer was operated on a full scan mode from 50 to 500 at an acquisition rate 20 spectra / sec. To provide a set of data that can be used to assess overall reproducibility and to correct for potential analytical variations, a pooled plasma sample containing aliquots from all study subjects (or representative subjects depending on the number of samples to be tested) were used as a study QC. The QC samples for this project were prepared with the test samples and were injected at regular intervals (after every 10 test samples for GC-TOFMS and after every 12 test samples for UPLC-QTOFMS, respectively) to allow evaluating overall process variability and monitoring platform performance. The acceptance criterion for RSD is typically set to < 20% (add citation/reference) or < 30% (add reference).

### Compound annotation and marker selection

All the features that pass the quality control are subject to compound annotation. This is performed using an in-house library containing ∼1,000 mammalian metabolites (reference standards). Commercial databases including NIST library 2011, LECO/Fiehn Metabolomics Library, etc., are also used for compound annotation and verification. For the LC-TQMS data, the data were collected with multiple reaction monitor (MRM), and the cone and collision energy for each metabolite used the optimized settings from QuanOptimize application manager (Waters) and the metabolites were all annotated with reference standards. For the GC−TOFMS generated data, identification was processed by comparing the mass fragments and the Rt with our in-house library or the mass fragments with NIST 05 Standard mass spectral databases in NIST MS search 2.0 (NIST, Gaithersburg, MD) software using a similarity of more than 70%.

### Metabolomics data pre-processing

The raw LC-MS/MS data files were processed with TargetLynx Application Manager (Waters Corp., Milford, MA) to extract peak area and retention time of each metabolite. The raw GC-TOFMS data files were processed with ChromaTOF software (Leco Corp., St Joseph, MI) to extract peak signal and retention times for each metabolite. The detected metabolites were annotated with our internal metabolite database using an automated mass spectral data processing software package, ADAP-GC ^27^.

### Metabolomics data processing

Samples were received in 3 different batches. Batch # 1 (N=36), #2 (N=21) and #3 (N=30) detected 93, 120 and 106 untargeted metabolites respectively. A total of 79 untargeted metabolites were common in the discovery cohort (batch 1 and 2) and were combined with the 151 targeted metabolites, yielding 230 metabolites total in the training set. 151 metabolites were identified from LC, and 79 metabolites were from GC. Power analysis was done using the module implemented in MetaboAnalystR^33^, on both the whole metabolite list and the selected 29 metabolites. We conducted data pre-processing similar to the previous report^34^. Briefly, we used K-Nearest Neighbors (KNN) method to impute missing (∼ 8%) metabolomics data ^35^. Using KNN method, metabolites missed value was predicted from the mean of the k nearest neighbors. To adjust for the offset between high and low-intensity features, and to reduce the heteroscedasticity, the logged value of each metabolite was centered by its mean and autoscaled by its standard deviation ^36^. We used quantile normalization to reduce sample-to-sample variation ^37^. We applied partial least squares discriminant analysis (PLS-DA) to visualize how well metabolites could differentiate the obese from normal samples. To explore the contribution of different clinical/physiological factors to metabolomics data, we conducted source of variation analysis. We used comBat Bioconductor R package ^38^ to adjust for the batch effects in the metabolomics data.

### Classification modeling and evaluation

To reduce the dimensionality of our data (230 metabolites vs 57 samples), we selected the unique metabolites associated with separating obese and normal status. To achieve this, we used a penalized logistic regression method called elastic net that was implemented in the glment R package ^39^. Elastic net method selects metabolites that have non-zero coefficients (y-axis, Figure 3S-C) at different value of lambda (x-axis, Figure 3S-C), guided by two penalty parameters alpha and lambda ^39^. Alpha sets the degree of mixing between lasso (when alpha=1) and the ridge regression (when alpha=0). Lambda controls the shrunk rate of coefficients regardless of the value of alpha. When lambda equals zero, no shrinkage is performed and the algorithm selects all the features. As lambda increases, the coefficients are shrunk more strongly and the algorithm retrieves all features with non-zero coefficients. To find optimal parameters, we performed 10-fold cross-validation for feature selection that yield the smallest prediction minimum square error (MSE), similar to previous studies^40^. We then used the metabolites selected by the elastic net to fit the regularized logistic regression model. Three parameters were tuned: cost, which controls the trade-off between regularization and correct classification, logistic loss and epsilon, which sets the tolerance of termination criterion for optimization.

**Figure 3:**
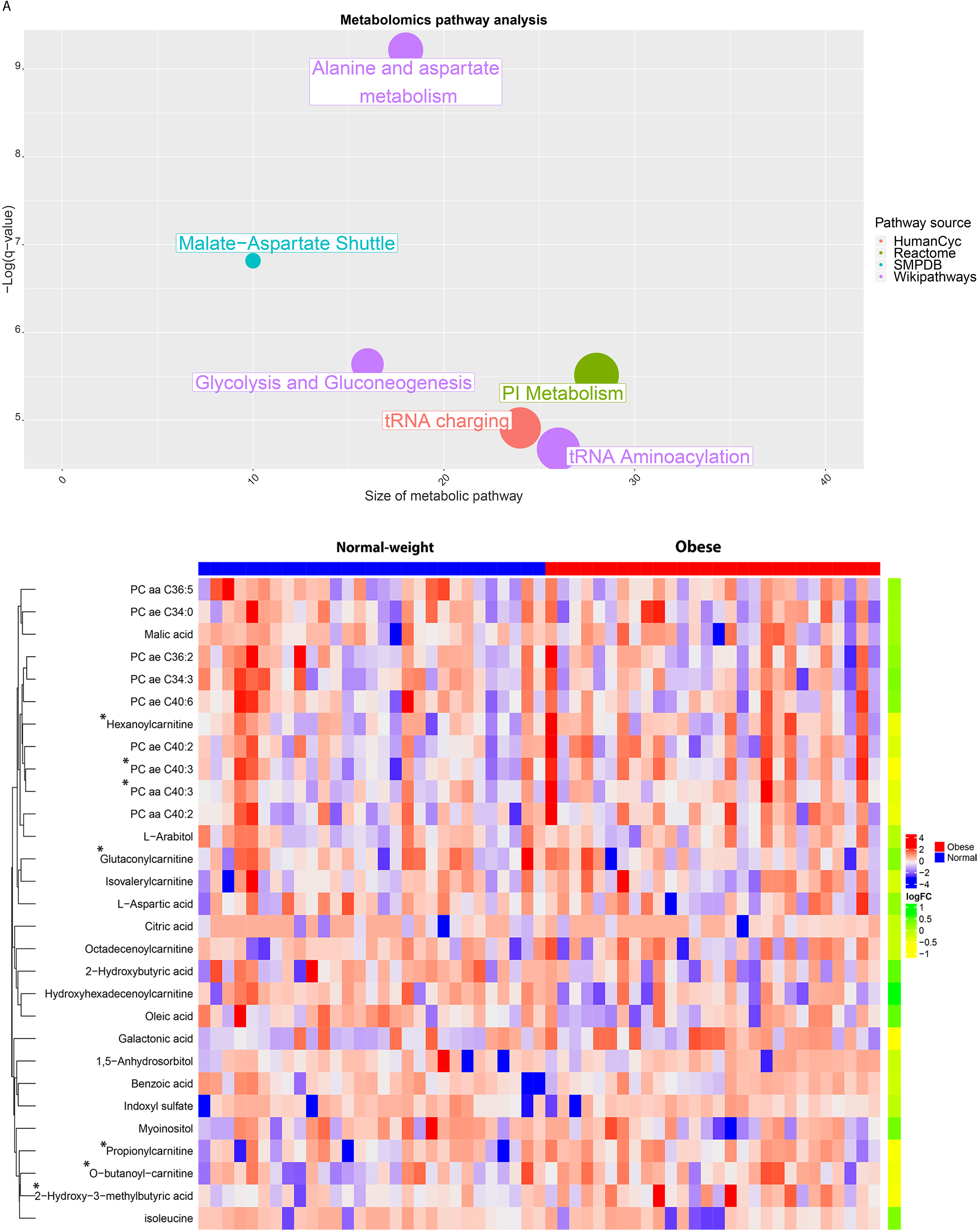
Analysis of the 29 selected metabolites. (A) Heatmap of selected metabolites separated by maternal group. * indicates metabolites that shows significant p-values (P<0.05, t-test) individually. (B) Pathway analysis of the 29 metabolites. X-axis shows size of metabolomic pathway. Y-axis shows the adjusted p-value calculated from CPDB tool. The size of the nodes represents the size of metabolomic pathway (number of metabolites involved in each pathway). The color of the nodes represents the source of these pathways.

To construct and evaluate the model, we conducted cross validation using five folds. We trained the model on four folds (80% of data) using leave one out cross validation (LOOCV) and measured model performance on the remaining fold (20% of data). We carried out the above training and testing five times on all folds’ combination. We plotted the receiver-operating characteristic (ROC) curve for all folds’ prediction using pROC R package. To adjust confounding other clinical covariants such as ethnicity, gravidity and parity, we reconstructed the metabolomics model above by including these factors.

### Metabolites importance score

To rank the metabolites based on their contribution to the model performance, we used the model-based approach implemented in the Classification And REgression Training (CARET) R package^41^. The importance of each metabolite is evaluated individually using a “filter” approach^42^. The ROC curve analysis is conducted on each metabolite to predict the class using a series of cutoffs. The sensitivity and specificity are computed for each cutoff and the ROC curve is computed. The trapezoidal rule is used to compute the area under the ROC curve. This area is used as the measure of variable importance. These scores were scaled to have a maximum of 100.

### Metabolomics pathway analysis

We performed metabolomic pathway analysis on metabolites chosen by the elastic net method using Consensus Pathway DataBase (CPDB). We used rcorr function implemented in Hmisc R package to compute the correlations among clinical and metabolomics data.

### Source of variation analysis

To estimate the relative contribution of the confounder factors such as maternal age and ethnicity to the variability of the metabolomics data, we built the ANOVA model using the metabolomics and cofounder factors, the resulting F-value is used to calculate p-values.

### Data availability

The metabolomics raw data files as well as processed data generated by this study have been deposited to Metabolomics workbench repository (https://www.metabolomicsworkbench.org/, Study ID ST001114).

The list of unknown metabolites and their characteristics are included in Supplementary Tables 1, 2 and 3.

## Results

### Cohort subject characteristics

Our discovery cohort consisted of 57 samples (29 normal and 28 obese subjects) from different ethnic groups. It consisted of three ethnic groups: Caucasian, Asian and Native Hawaiian and other Pacific Islander (NHPI). Women undergoing scheduled cesarean delivery were included based on the previously described inclusion and exclusion criteria (Methods). Demographical and clinical characteristics in obese and control groups are summarized in Table 1. In the Caucasian group (10 mothers), 6 were categorized as non-obese and 4 as obese. In the Asian group (23 mothers), 16 were categorized as non-obese and 7 as obese. In the NHPI group (24 mothers), 7 (24%) were categorized as non-obese and 17 (61%) as obese. The variation in recruitment of cases versus controls in each ethnic background reflects the demographics in Hawaii. Compared to mothers of normal pre-pregnant BMI, obese mothers had significantly higher pre-pregnancy BMI (33.51+/- 4.49 vs 21.89 +/- 1.86 kg/m2, p=9.18e-11). Mothers had no statistical difference regarding their ages (32.10 +/- 4.88 vs 32.48 +/- 5.66, p=0.7) or gestational age (39.04 weeks+/- 0.22 vs 38.93 +/-0.45 p=0.38), excluding the possibility of confouding from these factors. Babies of obese mothers had significantly (P=0.03) higher birth weight averages in comparison to the normal pre pregnant weight group, consistent with earlier observations ^43, 44^.

**Table 1:** Demographical and clinical characteristics in obese and control groups.

|  | Control(n=29) | Case(n=28) | P-value* | Confounder** | SOV*** |
| --- | --- | --- | --- | --- | --- |
|  | Mean (SD) |  |  |  |  |
| Maternal age, years | 32.48 (5.66) | 32.10 (4.88) | 0.78 | Yes | 0.79 |
| Paternal age, years | 34.68(7.14) | 35.21(6.43) | 0.79 | Yes | 0.94 |
| Pre-pregnancy BMI, kg/m2 | 21.89(1.86) | 33.51(4.49) | 1.12 e-14 | Yes | 1.61 |
| Gestational Age, Weeks | 39.04(0.218) | 38.93(0.45) | 0.3812 | Not included |  |
| Net weight gain | 30.85(10.92) | 29.4(13.55) | 0.7335 |  |  |
| Baby weight (kg) | 3.29(0.32) | 3.54(0.5) | 0.03 |  |  |
| Head Circle (cm) | 34.89(1.10) | 35.55(1.36) | 0.05 |  |  |
| Baby length (cm) | 51.3(1.9) | 51.4(2.36) | 0.8 |  |  |
| **Parity** |  |  | 0.03 | Yes | 0.80 |
| 0 | 5 | 2 |  |  |  |
| 1 | 16 | 7 |  |  |  |
| 2 | 7 | 10 |  |  |  |
| 3 and above | 1 | 9 |  |  |  |
| **Gravidity** |  |  | 0.12 | Yes | 0.98 |
| 1 | 5 | 1 |  |  |  |
| 2 | 12 | 5 |  |  |  |
| 3 | 7 | 8 |  |  |  |
| 4 and above | 5 | 14 |  |  |  |
| **Maternal Ethnicity** |  |  | 0.01 | Yes | 0.73 |
| Caucasian | 6 | 4 |  |  |  |
| Asian | 16 | 7 |  |  |  |
| Pacific island | 7 | 17 |  |  |  |
| **Paternal Ethnicity** |  |  | 0.03 | Yes | 0.88 |
| Caucasian | 8 | 3 |  |  |  |
| Asian | 14 | 9 |  |  |  |
| Pacific island | 7 | 16 |  |  |  |

\*Categorical variables were compared using chi-square test, whereas continuous variables were compared using *t* test.
\*\* Yes if the factor is accounted for when building the classifier model
\*\*\* The source of variation of the factor. It represents the contribution of the factor to the variability of the metabolomics data. It is the F-stats that were taken from a linear regression ANOVA model comprising maternal and paternal age and ethnicity, parity and gravidity (Figure 1A).

### Preliminary assessment of metabolomics results

Our discovery cohort has a total of 230 metabolites, including 79 untargeted and 151 targeted metabolites. To explore which clinical or physiological covariates were associated with the variations in the metabolomics, we conducted source of variation analysis using a linear mixed model that includes multiple clinical variables. Figure 1A shows the F value for each factor in the ANOVA model relative to the error. The Y-axis is the F value and X-axis are the factors specified in ANOVA model and the Error term. Indeed, maternal obesity was the predominantly the most important factor contributing to metabolomics difference, rather than other factors (Fig 1A). To test if these metabolites allow a clear separation between the obese and normal-weight subjects, we used elastic net regularization based logistic regression, rather than the partial least squares discriminant analysis (PLS-DA) model, a routine supervised multivariate method which only yielded modest accuracy AUC=0.62 (Fig 2S-B). PLS-DA has high probability to develop models that fit the training data well, however, produce poor predictive performance on the validation set^45^.

**Figure 1:**
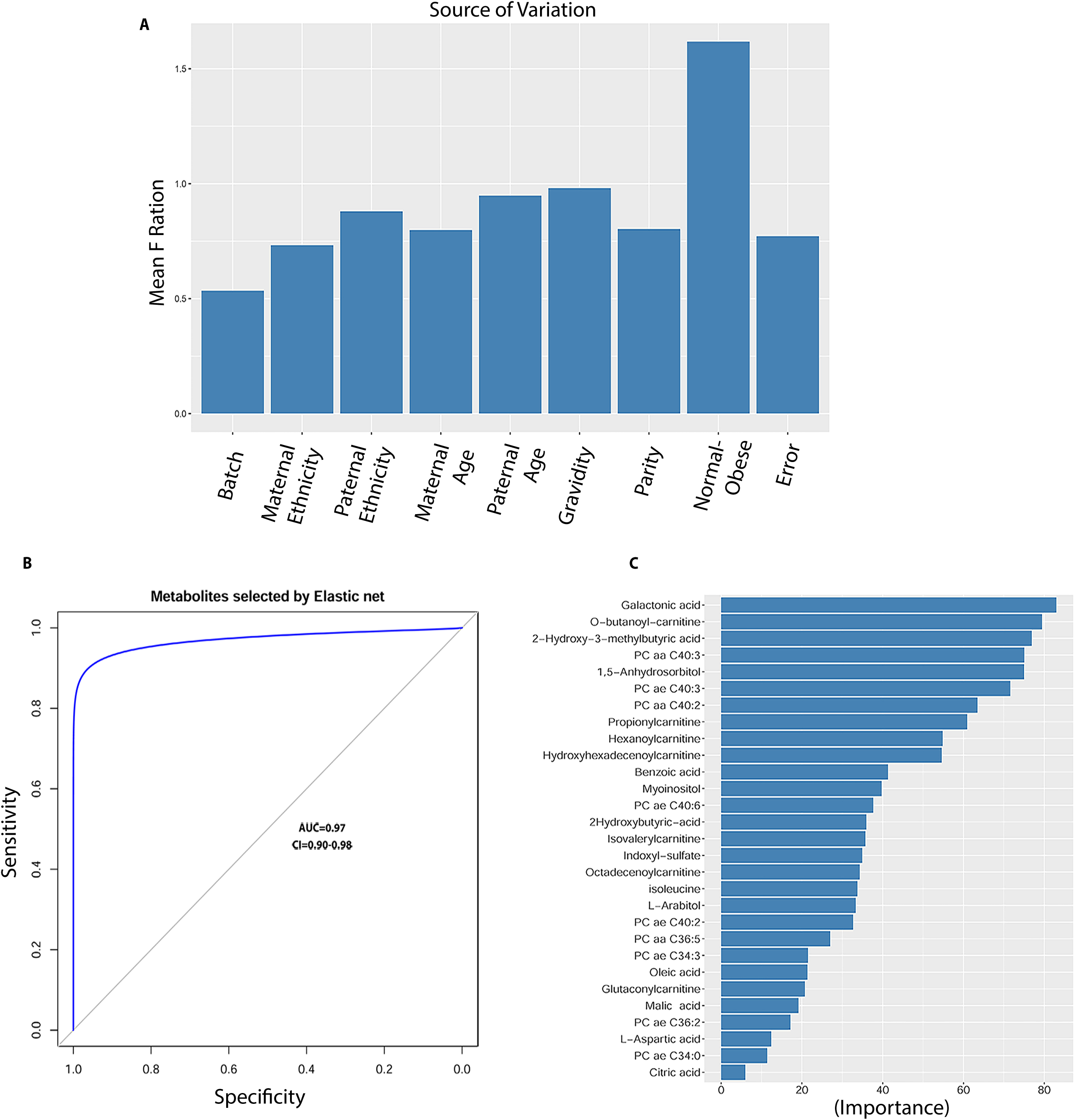
Source of variation and accuracies of logistic regression models and important features selected by the metabolomics model. (A) ANOVA plot of clinical factors using the metabolites levels in cord blood samples. Averaged ANOVA F-statistics are calculated for potential confounding factors, including obesity, gravida, parity, paternal and maternal age and ethnicity. (B) Model accuracy represented by classification Receiver Operator Curves (ROCs). (C) The ranking of contributions (percentage) of selected metabolomics features in the model.

**Figure 2:**
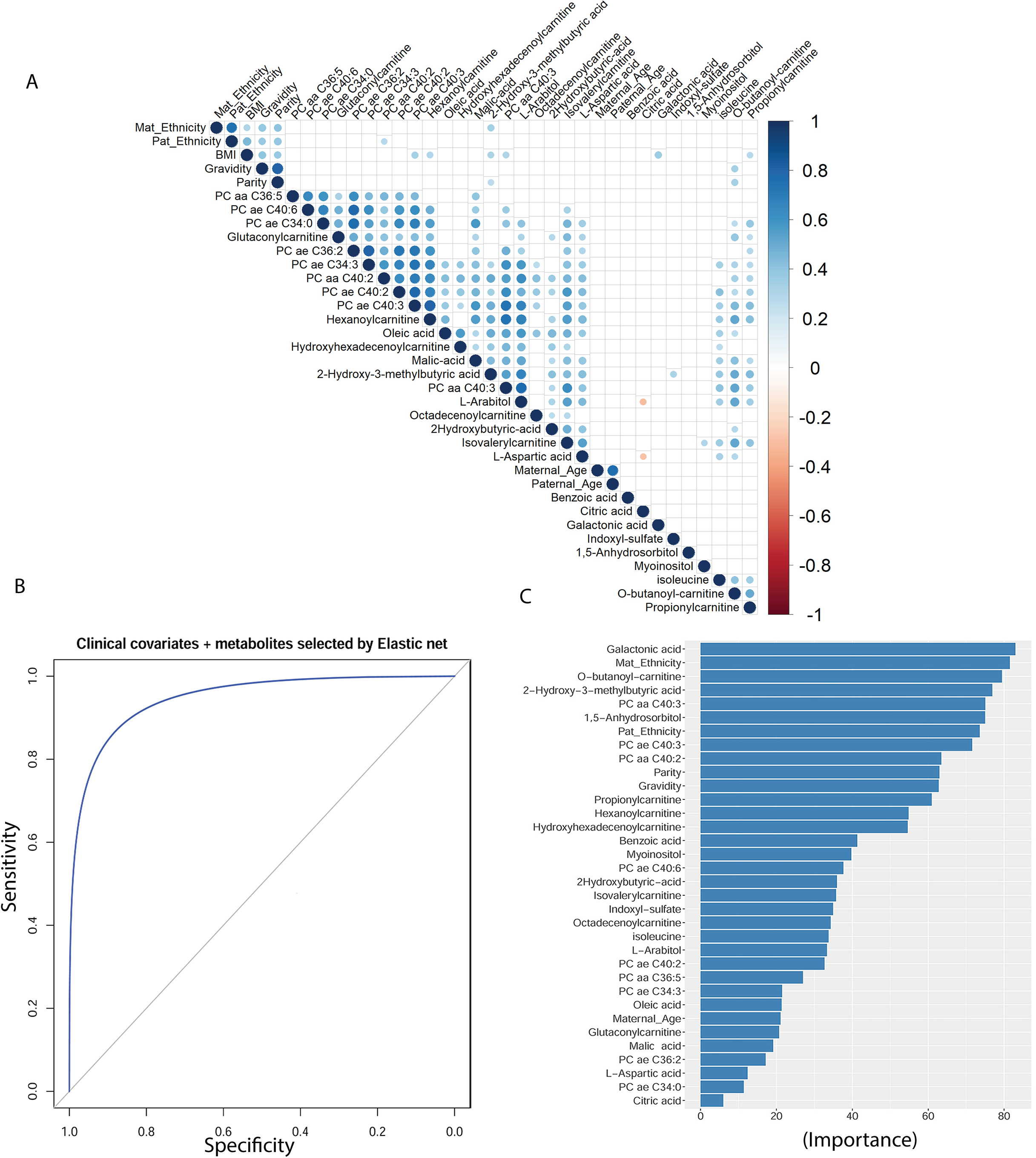
(A) Correlation coefficients among demographical/physiological factors and the metabolomics data. Blue color indicates positive correlations and red indicated negative correlations. (B) Receiver Operator Curves (ROCs) of the combined model with metabolomics and physiological/demographic data. (C). The ranking of contributions (percentage) of selected features in the model (B).

On the other hand, elastic net offers potential improvements over PLS-DA due to the presence of the constrain, which promotes sparse solutions. Moreover, elastic net regularization overcomes the limitation of either ridge and lasso regularization alone and combines their strengths to identify optimized set metabolites [25]. Using the optimized regularization parameters (Fig. 3S), we identified a total of 29 metabolite features (Table 2), which together yields the highest predictive performance with AUC=0.97, 95 % CI=[0.904-0.986] in 20% hold-out test dataset (Figure 1B). The 29 selected metabolites by the elastic-net collectively yields a statistical power of 0.93, much improved from the initial power of 0.67 estimated from the total 230 metabolites (Fig. 4S), and supports the validity of the statistical modeling approach. Among them, six metabolites have large contributions to the separations between the case and controls, with an importance score of at least 70% individually (Figure 1C). These are galactonic acid, butenylcarnitine (C4:1), 2-hydroxy-3-methylbutyric acid, phosphatidylcholine diacyl C40:3 (PC aa C40:3), 1,5-anhydrosorbitol, and phosphatidylcholine acyl-alkyl 40:3 (PC ae C40:3). Thus, metabolites selected by the elastic net method indeed improved the prediction power of the model overall.

**Table 2:**
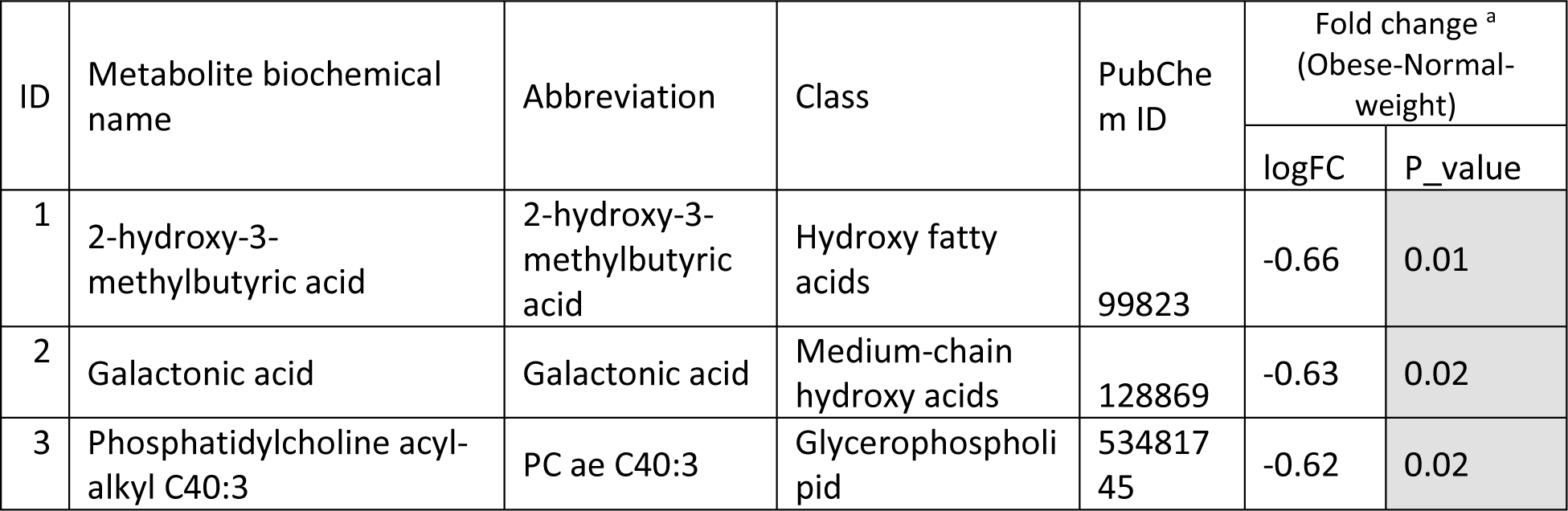
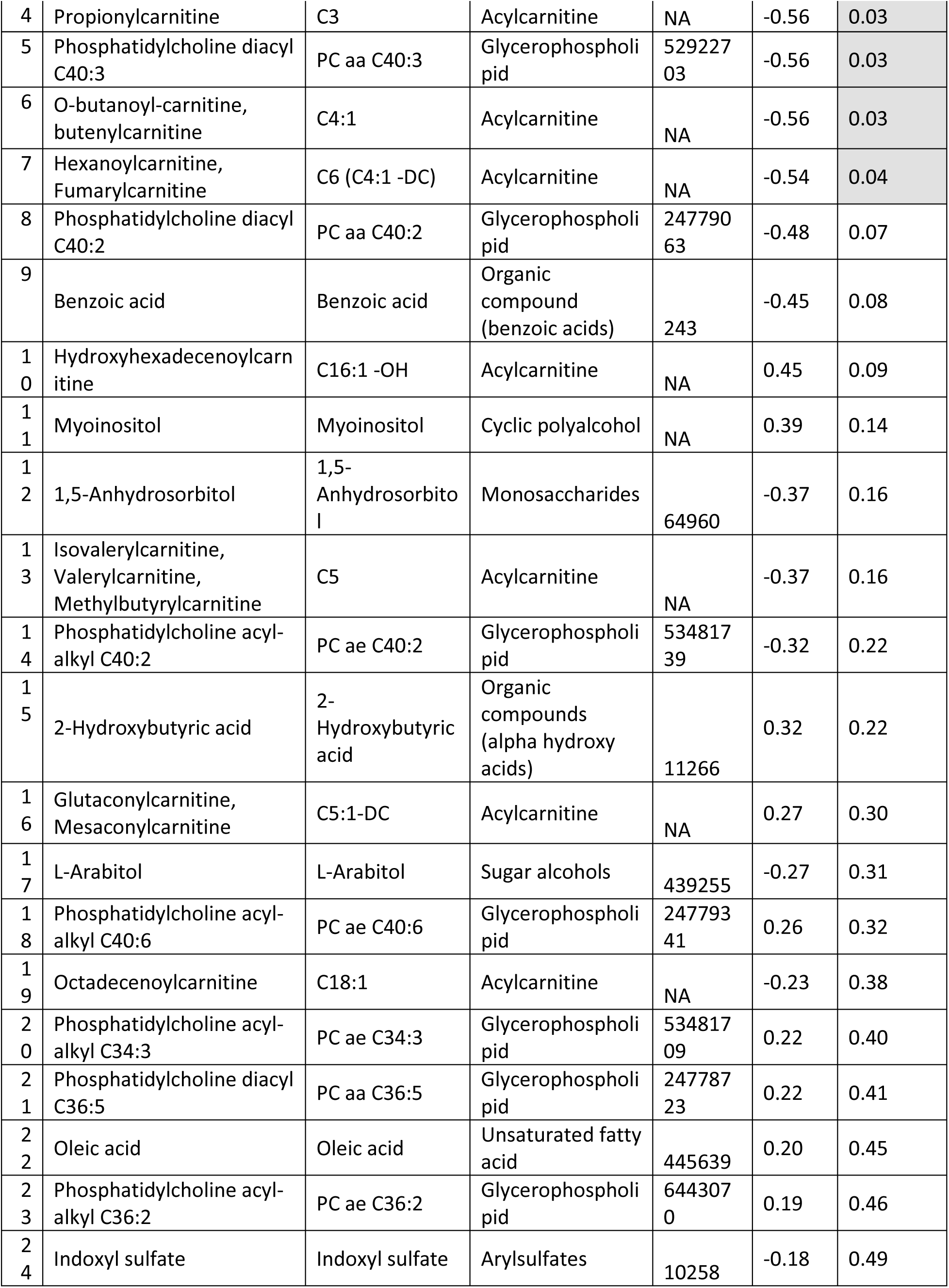
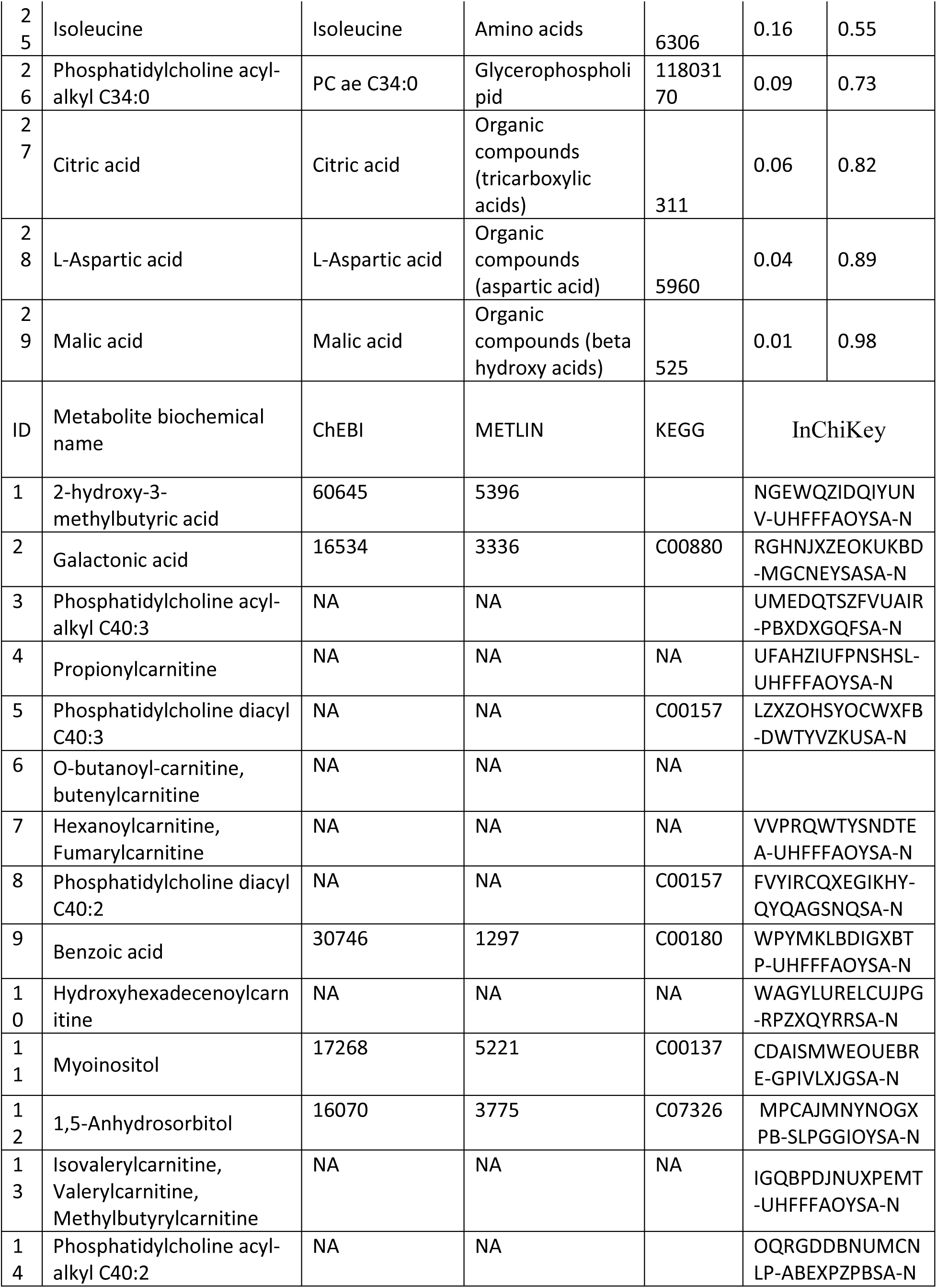
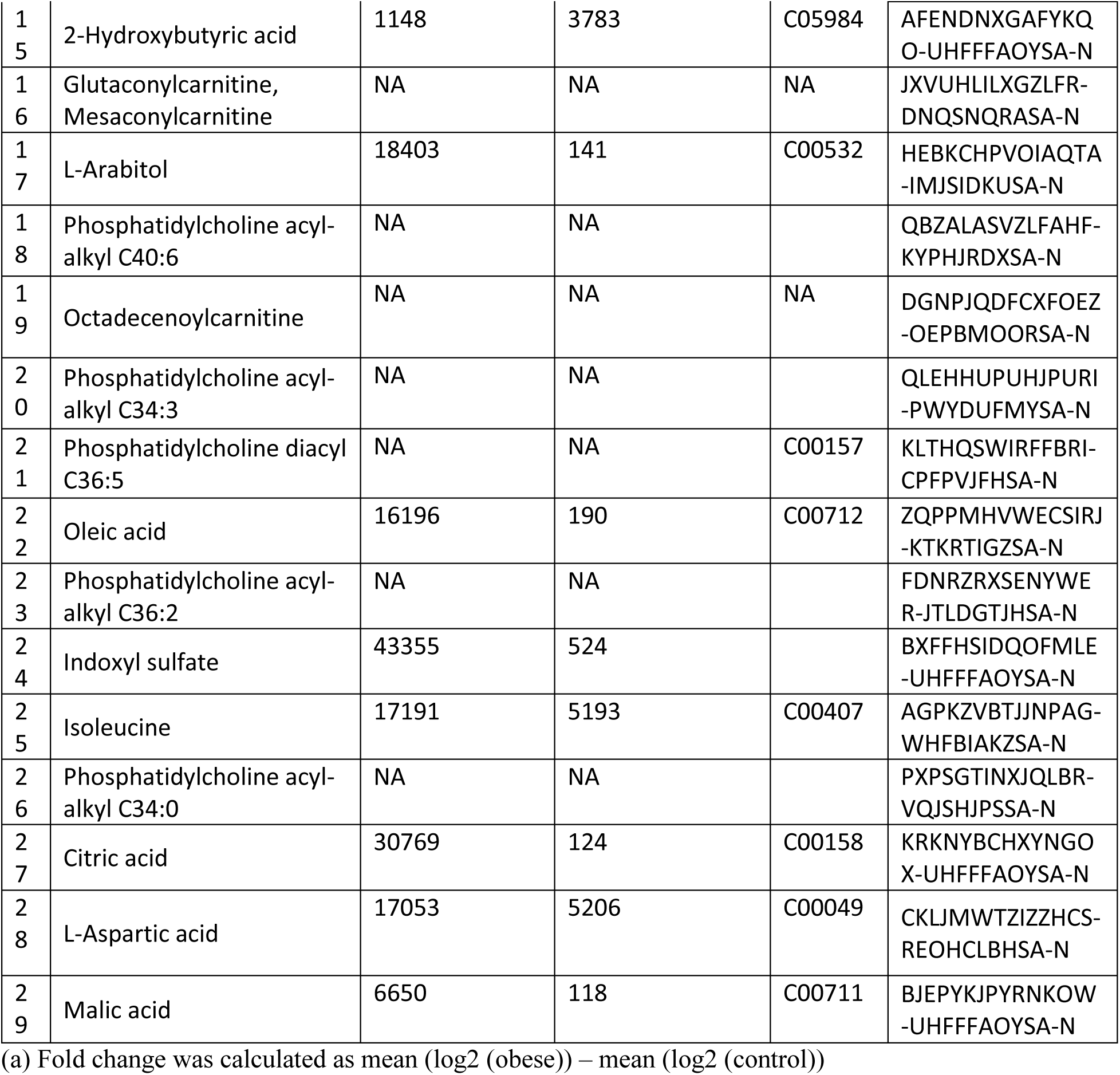
A list of metabolites associated with obese-control maternal status, selected by elastic net regularization based logistic regression. The metabolites are sorted by the average log fold change of cases over controls.

**Figure 4:**
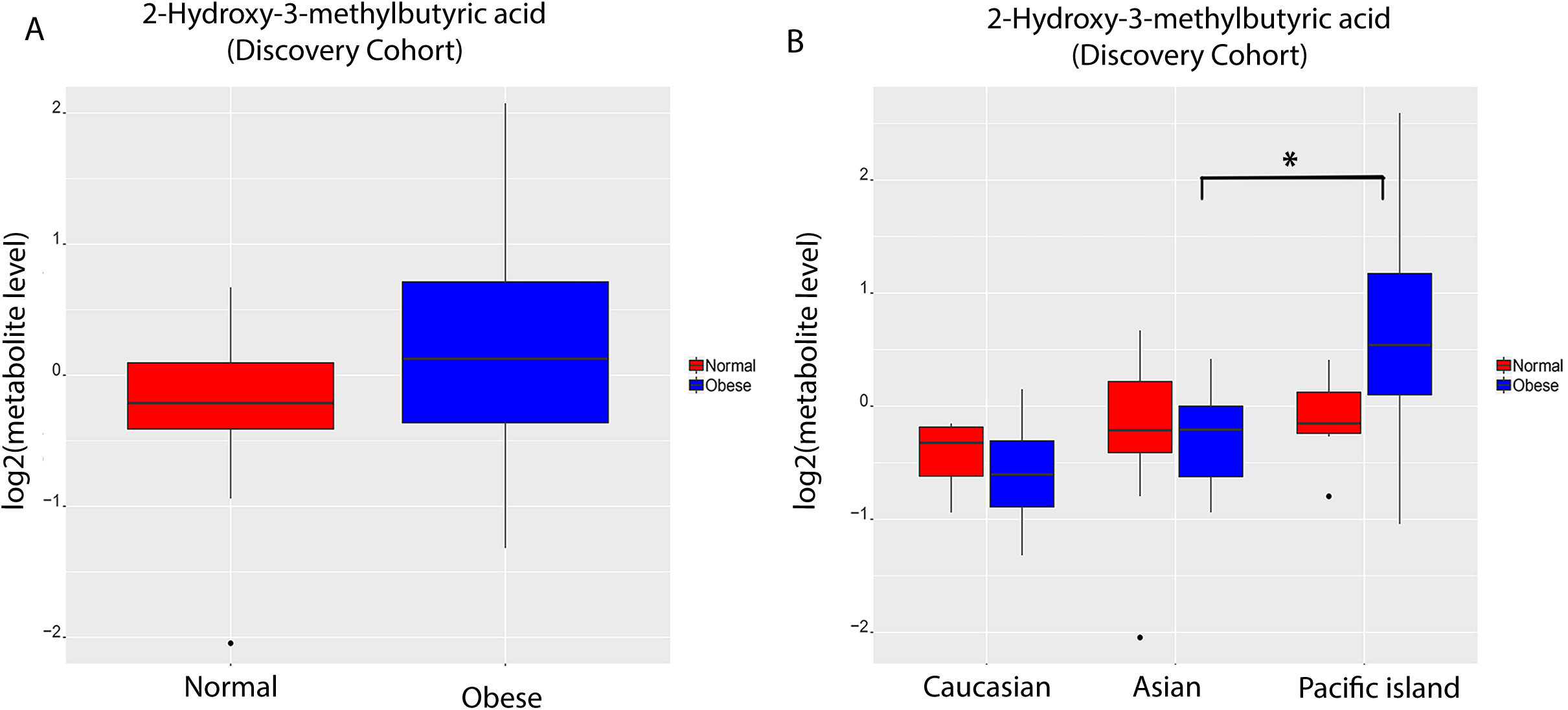
Box plot of 2-hydroxy-3-methylbutyric acid in the discovery cohort, stratified by (**A)** normal (n=29) and obese (n=28) subjects, and further by (**B)** the three ethnic groups: Asian, Caucasian and Native Hawaiian and Pacific Islanders (NHPI).

### Calibrated maternal-obese predictive model with consideration of confounding

For statiscal rigor, it is important to consider possible clinical confounders (if available), such as maternal and paternal ethnicity and parity (Table 1) that we collected for additional calibration. Towards this, we conducted two investigations. First, we explored the correlations among the demographic factors and metabolomics data. It is evident that several metabolites are correlated with maternal and paternal ethnicity, gravidity, and/or parity (Figure 2-A). For example, maternal ethnicity is positively correlated with 2-hydroxy-3-methylbutyric acid. Secondly, we built a logistic regression model using the above-mentioned four covariates alone (parity, gravidity, maternal and paternal ethnicity). This model yields a modest AUC of 0.701 95% CI= [0.55-0.82] (Figure 4S-A), again suggesting existence of confounding. These observations prompted us to recalibrate the 29-metabolite elastic net model, by adjusting the metabolomics model using maternal and parental age and ethnicity, gravidity and parity (Figure 2B). The resulting modified model remains to have very high accuracy, with AUC= 0.947, 95% CI= [0.88-0.98]. In the new model, besides the original 6 metabolite features, maternal ethnicity and paternal ethnicity also have importance scores greater than 70% (Figure 2C).

### Metabolite features and their pathway enrichment analysis

The 29 metabolite features selected by the model belong to acylcarnitine, glycerophospholipid, amino acids and organic acids classes. Their log fold changes ranged from −0.66 (2-hydroxy-3-methylbutyric acid) to −0.45 (hydroxyhexadecenoylcarnitine, or C16:1-OH) (Figure 3A and Table 2). Among them, 15 metabolites were higher in obese associated cord blood samples, including 2-hydroxy-3-methylbutyric acid, galactonic acid, PC ae C40:3, propionylcarnitine (C3), PC aa C40:3, o-butanoyl-carnitine (C4:1), hexanoylcarnitine (C6 (C4:1 -DC)), ohosphatidylcholine diacyl C40:2 (PC aa C40:2), benzoic acid, 1,5-anhydrosorbitol, isovalerylcarnitine (C5), PC ae C40:2, L-arabitol, octadecenoylcarnitine (C18:1) (Figure 3A, Table 2). The remaining 14 metabolites are lower in obese associated cord blood samples: malic acid, aspartate, citric acid, PC ae C34:0, isoleucine, PC ae C36:2, oleic acid, PC aa C36:5, PC ae C34:3, PC ae C40:6, C5:1-DC, 2-hydroxybutyric acid, myoinositol, and C16:1 -OH (Figure 3A, Table 2). The individual metabolite levels of hexanoylcarnitine (C6(C4:1-DC)), o-butanoyl-carnitine (C4:1), PC aa C40:3, propionylcarnitine (C3), PC ae C40:3, galactonic acid, and 2-hydroxy-3-methylbutyric acid increased significantly in obese cases (p<0.05, t-test).

To elucidate the biological processes in newborns that may be effected by maternal obesity, we performed pathway enrichment analysis on the 29 metabolite features, using Consensus pathway database (CPDB) tool ^46^. We combined multiple pathway databases including KEGG, Wikipathways, Reactome, EHNM and SMPDB. A list of the enriched pathways is plotted with adjusted p-value q<0.05 (Figure 3B). We removed large pathways such as SLC-mediated transmembrane transport for non-specificity. Among the filtered pathways, alanine and aspartate metabolism is the most significantly enriched pathway (q=0.004). One should note that given the small number of identified metabolites in the dataset, the pathway analysis may have limited reliability.

### The influence of ethinicity on metabolite levels

In general, the level of 2-hydroxy-3-methylbutyric acid in obese subjects is higher than compared to the normal-weighted subjects (Figure 4A). Our earlier correlational analysis suggested that maternal ethnicity may be correlated with 2-hydroxy-3-methylbutyric acid level (Figure 2A). To confirm this, we conducted 2-way ANOVA statistical tests and indeed obtained significant p-value (P=0.023, chi-square test). We thus stratified the levels of 2-hydroxy-3-methylbutyric acid by ethnicity (Figure 4B). There was no significant difference in normal pre pregnant-weight subjects across the three ethnic groups (Figure 4B). However, in cord blood samples associated with obese mothers, the concentration of 2-hydroxy-3-methylbutyric acid was much higher in NHPI, as compared to Caucasians (p=0.05) or Asians (p=0.04) (Figure 4B). 2-hydroxy-3-methylbutyric acid originates mainly from ketogenesis through the metabolism of valine, leucine and isoleucine ^47^. Since all subjects fasted 8 hours before the C-section, we expect the confounding from diets is minimized among the three ethnic groups. Thus the higher 2-hydroxy-3-methylbutyric acid level may indicate the higher efficiency of ketogenesis in babies born from obese NHPI mothers.

### Validation on an independent cohort

We subsequently collected a new set of 30 patients (18 normal-weight and 12 obese). We decided to treat this set as “validation cohort”, following the convention of machine-learning dataset design, as samples were processed in different times/batches. We aimed to test if the previous model built on the 57 samples is predictive, given the modest size and heterogeneity among samples. We then performed new metabolomics measurements and processed the data as earlier described. The model built on 57 samples yields an AUC of 0.822 (95% CI= [0.74-0.89], Figure 5A) in the new set of 30 samples, confirming the reproducibility of our findings. Moreover, we observed a similar trend of higher concentration of 2-hydroxy-3-methylbutyric acid in the obese subjects compared to normal-weight (Figure 5B). Importantly, the levels of 2-hydroxy-3-methylbutyric acid has a similar trend in NHPI compared to Asians and Caucasians (p=0.001) in the obese group, whereas no statistical difference between ethnicities exists in the control group (Figure 5C). Moreover, within this cohort, four of the six metabolites that had large contributions to the separations between case/control (importance score > 70%) in the discovery cohort, had consistent trend of changes in the validation cohort.

**Figure 5:**
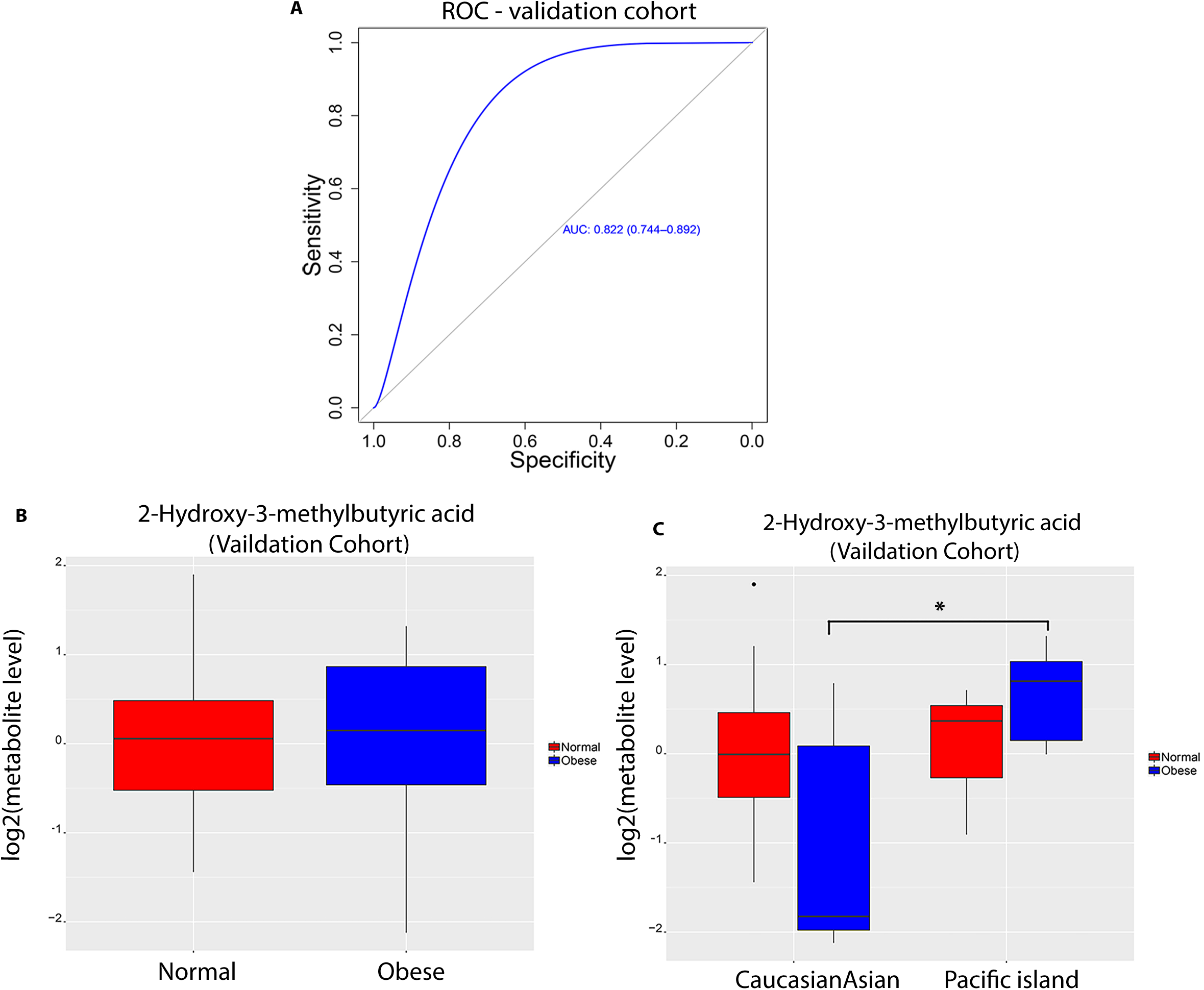
Validation with a subsequent cohort (n=30). (**A)** Accuracy on classifying cases vs controls in the validation cohort, using the model built on the discovery cohort as shown in Fig 2(B). **(B-C)** Box plots of 2-hydroxy-3-methylbutyric acid, stratified by (**B)** normal (n=18) and obese (n=12) subjects, and further by (**C)** ethnic groups of Asians/Caucasians vs. Native Hawaiian and Pacific Islanders (NHPI). Asians (n=2) and Caucasians (n=3) were combined, as the number of patients of these ethnicities in the obese group is small in the obese group. *: statistically significant with p-value <0.05 (t-test).

## Discussion

This study aims to distinguish key cord blood metabolites associated with maternal pre-pregnancy obesity. As maternal obesity is a health condition rather than a disease, we had to set stringent inclusion and exclusion crtieria to exclude as many confouding factors as possible, to ensure the quality of the metabolomics data. To avoid the sources of confounding from labor and vaginal delivery (diets, multiple operators due to unpredictable delivery time etc.), we only targeted mothers having elective C-sections. We also excluded obese mothers who had known complications during pregnancy, such as pre-gestational diabetes, smoking, and hypertension. To minimize confounding due to maternal diet, all subjects fasted 8 hours before the Cesarean section. These criteria helped to improve the quality of the samples and hence metabolomics data, albeit the size of the study is modest.

Such careful experimental design did yield good data quality, as the source of variation analysis did show that maternal obesity is the only dominate factor contributing to metabolomics difference in the cord blood. Additionally, we conducted rigorous statistical modeling and found that metabolites can distinguish the two maternal groups with accuracy as high as AUC=0.97 under cross-validation (or 0.947 after adjusting for confounding effects). Among all metabolites and physiological/demographic features selected by the combined model, galactonic acid has the largest impact on the model performance (importance score =86%). Galactonic acid is a sugar acid and breakdown product of galactose. When present in sufficiently high levels, galactonic acid can act as an acidogen and a metabotoxin which has multiple adverse effects on many organ systems. Galactonic acid, was previously shown to be associated with diabetes in a mouse model, due to a proposed mechanism of oxidative stress ^48^. On the other hand, maternal ethnicity has the largest impact among physiological factors (importance score =84%).

Few cord blood metabolomics studies have been carried out to associate with maternal obesity directly, or birth weight ^23, 49, 50^. In a recent Hyperglycemia and Adverse Pregnancy Outcome (HAPO) Study, Lowe et al. reported that branched-chain amino acids and their metabilites, such as valine, phenylalanie, leucine/isoleucine and AC C4, AC C3, AC C5 are associated with maternal BMI in a meta-analysis over 4 large cohorts (400 subjects in each) ^50^. In another study to associate cord blood metabolomics with low birth weight (LBW), Ivorra et al. found that newborns of LBW (birth weight < 10th percentile, n = 20) had higher levels of phenylalanine and citrulline, compared to the control newborns (birth weight between the 75th-90th percentiles, n = 30) ^23^. They also found lower levels of choline, proline, glutamine, alanine and glucose in new borns of LBW, however, there was no significant differences between the mothers of the two groups. In our study, isoleucine is also identified as one of the 29 metablite features related to maternal obesity; although alanine iteself is not selected by the model to be a maternal obesity biomarker in cord blood, we did find that alanine and aspartate metabolism are enriched in the cord blood samples associated with maternal obesity group.

Metabolomics pathway analysis on the metabolite features in the model identified six filtered significant pathways (Figure 3A). Among them, alanine and aspartate metabolism was previously reported to be associated with obesity ^51 52^. Aspartate and alanine cycling has known association with insulin resistance and metabolic related diseases, such as cancer ^53, 54^. Alanine, a highly gluconeogenic amino acid, contributes to the development of glucose intolerance in obesity, as circulating alanine levels are elevated in obese mothers. Our study also demonstrates that in infants of obese mothers this pathway is also enriched. Additionally, glycolysis is the metabolic pathway that converts glucose into pyruvate, while gluconeogenesis is the reverse generating glucose from non-carbohydrate carbon substrates. The offspring of obese, but not normal-weight mothers in another study demonstrated downregulation of the glycolysis pathway (p=0.049) ^55^. Recent research showed that increase in hepatic gluconeogenesis was a major source of the total maternal glucose used by the fetus ^56^. Interestingly, 1,5-anhydrosorbitol, which has been shown to be a maternal marker of short-term glycemic control, was observed in our cord blood study as a marker too, likely from maternal origin. Thus, the changes in glycolysis and gluconeogenesis may suggest that obese mothers have greater glucose metabolism compared to normal controls. Phosphatidylinositol (PI) metabolism is a key regulator for energy metabolism. We found elevated levels of lipids such as PC aa 40:3 and PC ae 40:3 in obese subjects, in concert with this pathway. Altogether, the cord blood in babies of obese mothers demonstrates pathways enriched in metabolic syndrome and obesity, even though the phenotypic differences (obesity) does not exist in the babies but only mothers.

Notably, our study has identified 5 metabolites which are previously not reported in the literature with association to obesity or maternal obesity: galactonic acid, L-arabitol, indoxyl sulfate, 2-hydroxy-3-methylbutyric acid and citric acid. Except citric acid, all the other four metabolites are increased in obese associated cord blood samples. 2-hydroxy-3-methylbutyric acid concentrations varied by ethnicity, but only in babies born from obese pre-pregnant mothers. 2-hydroxy-3-methylbutyric acid is known to accumulate in high levels during ketoacidosis and fatty acid breakdown. Therefore, the higher elevation of 2-hydroxy-3-methylbutyric acid is likely due to increased cellular ketoacidosis and fatty acid breakdown in newborns from obese pre-pregnant mothers. To the best of our knowledge, this is the first study that shows differences in the 2-hydroxy-3-methylbutyric acid concentration levels among different ethnicities. Additionally, indoxyl sulfate is a metabolite of the amino acid tryptophan. As tryptophan is commonly found in fatty food, red meat and cheese, it is possible that high levels of indoxyl sulfate detected in the cord blood associated with obese pre-pregnant mothers could be due to the maternal high fat diet. Oppositely, citric acid, a compound associated with the citric acid cycle ^57^, is decreased in the cord blood associated with obese pre-pregnant mothers. This could be related to the lower vegetable and fruit consumptions among obese pre-pregnant mothers. In all, the data suggest that maternal obesity may impact offspring cord blood metabolites. Further research into the specific mode of action of these metabolites would be beneficial in understanding its association with maternal obesity.

This study may benefit from improvement in the future follow-up. We determined the subjects’ ethnicity by self-reporting rather than genotyping, due to the restriction of the currently approved IRB protocol. Additionally, there has been debate on the use of BMI as an indicator of obesity ^58^, more direct measures of body fat could be considered such as skin-fold thickness measurements, bioelectrical impedance and energy x-ray absorptiometry ^59, 60^. Moreover, dietary and exposomic data will be very interesting to study in a follow-up large-scale cohort with IRB approval. Nevertheless, this study has established relationships between cord blood metabolomics with maternal pre-pregnant obesity, which in turn is associated with socioeconomic disparities.

## Conclusion

In this study, we identified 29 cord blood metabolites that are associated with maternal obesity with high accuracy, in a discovery set of 57 samples and a validation set of 30 samples. These metabolites may have the potential to be maternal obesity-related bio-markers in newborns.

## Author Contributions

LXG envisioned the project, obtained funding, designed and supervised the project and data analysis. RJS, IC, PAB and SJC collected the samples. AG prepared the plasma samples. FMA analyzed the data. GX performed the metabolomics experiments. RJS, FMA, PAB, AG, GX, SJC and LXG wrote the manuscript. All authors have read, revised, and approved the manuscript.

## Acknowledgements

We thank Drs. Joseph Kaholokula and Alika Maunakea from the Native Hawaiian Health Department of University of Hawaii for giving suggestions. The authors acknowledge the services provided by the Molecular and Cellular Immunology Core which is funded in part by P30GM114737 from the Centers of Biomedical Research Excellence (COBRE) program of the National Institute of General Medical Sciences, a component of the National Institutes of Health. Dr. Lana Garmire’s research is supported by grants K01ES025434 awarded by NIEHS through funds provided by the trans-NIH Big Data to Knowledge (BD2K) initiative (http://datascience.nih.gov/bd2k), P20 COBRE GM103457 awarded by NIH/NIGMS, R01 LM012373 awarded by NLM, and R01 HD084633 awarded by NICHD to LX Garmire. Funding was also provided in part by the Department of Obstetrics and Gynecology, University of Hawaii. The metabolomics services were provided by the UH Cancer Center Metabolomics Shared Resource.

## Supplementary Information

### The following supporting information is available free of charge at ACS website http://pubs.acs.org

**Figure S1:** Discrimination of obese and normal groups by Partial Least Squares (PLS) method. **(A)** Discriminant analysis score plot for obese cases (Green) and normal (Red). (**B)** The accuracy of the 10 fold cross-validation of the PLS-DA model. R2 is the sum of squares captured by the model; Q2 is the cross-validation of R2.

**Figure S2:** Selection of metabolites using elastic net regularization. **(A)** Tuning alpha parameter, the parameter representing the degree of mixing between lasso (alpha=1) and the ridge regularization(alpha =0). Y-axis is the root mean square error of the 10-fold cross-validation. We selected alpha =0.22 as it gave us the minimum error. **(B)** Tuning lambda, the parameter controlling the shrunk rate of coefficients in the linear model. Y-axis is the misclassification error of the 10-fold cross validation. X-axis is the range of lambda, with the optimal lambda=0.008 as it gave us the minimum misclassification error. **(C)** The shrinkage coefficients of the metabolites using tuned alpha and lambda. Only metabolites with non-zero coefficient were be selected.

**Figure S3**: Accuracies of logistic regression models and important features selected by the clinical model. (A) Model accuracy represented by classification Receiver Operator Curves (ROCs). (B) The ranking of contributions (percentage) of selected clinical features in the model.

**Figure S4:** Power analysis and sample size estimation plot using (A) 230 metabolites and (B) 29 metabolites that were selected by elastic-net model.

**Table S1:** Unknown metabolites from batch #1

**Table S2:** Unknown metabolites from batch #2

**Table S3:** Unknown metabolites from batch #3

---

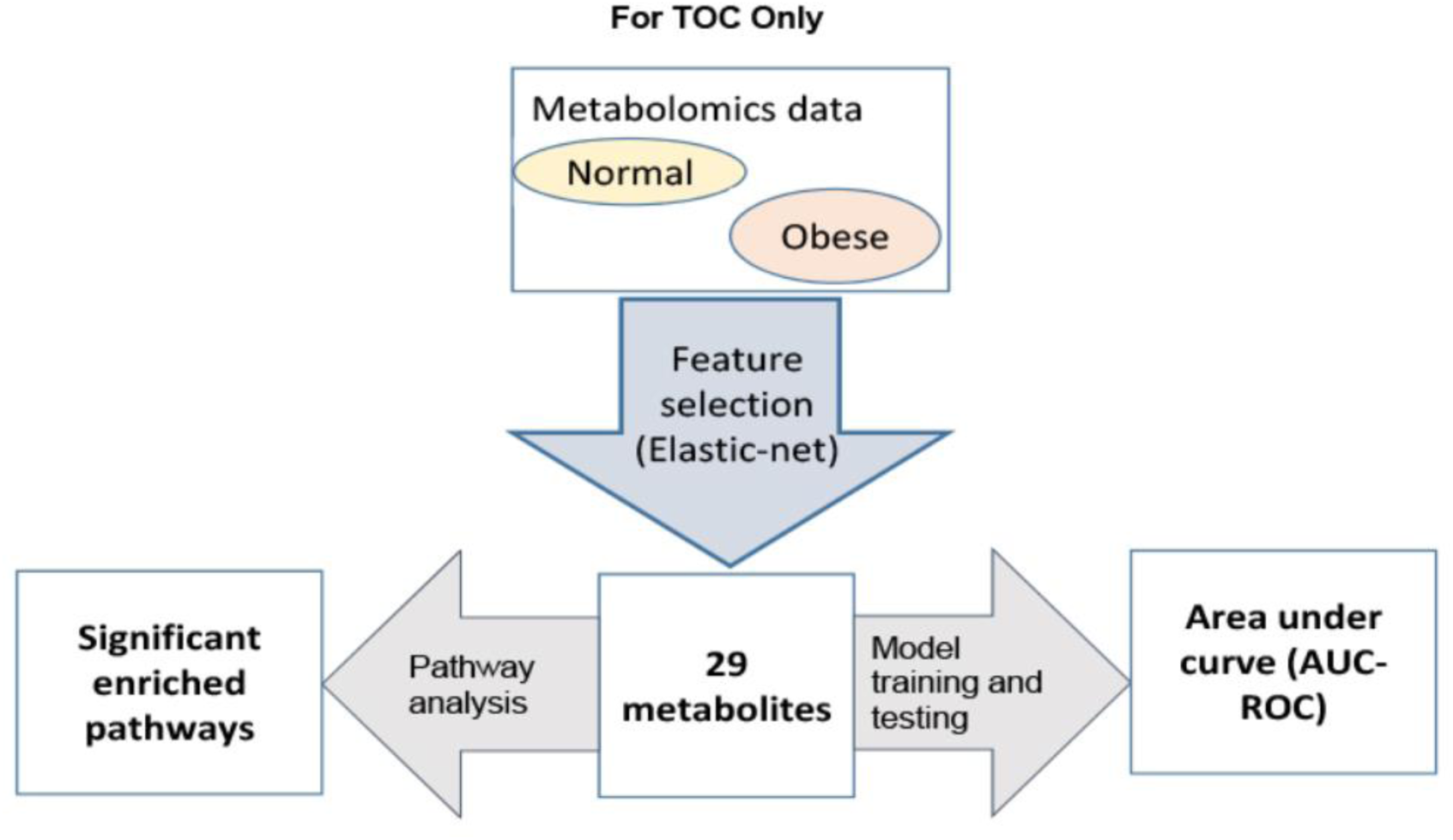

