## Supplementary material for "Pre-pregnant obesity of mothers in a multi-ethnic cohort is associated with cord blood metabolomic changes in offspring": Figure S1

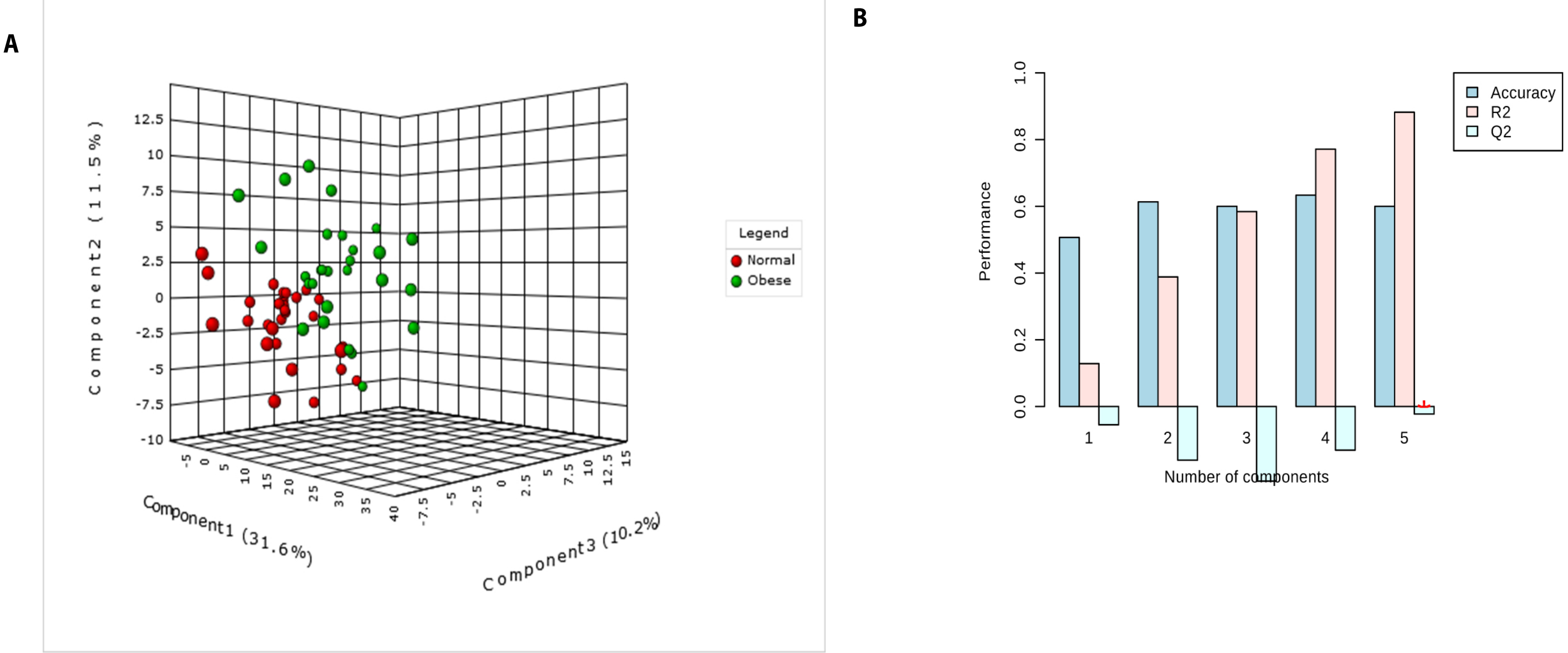


Supplementary Figure 1: Discrimination of obese and normal groups by Partial Least Squares (PLS) method. (A) Discriminant analysis score plot for obese cases (Green) and normal (Red). (B) The accuracy of the 10 fold cross-validation of the PLS-DA model. R2 is the sum of squares captured by the model; Q2 is the cross-validation of R2.
