## Supplementary material for "Pre-pregnant obesity of mothers in a multi-ethnic cohort is associated with cord blood metabolomic changes in offspring": Figure S2

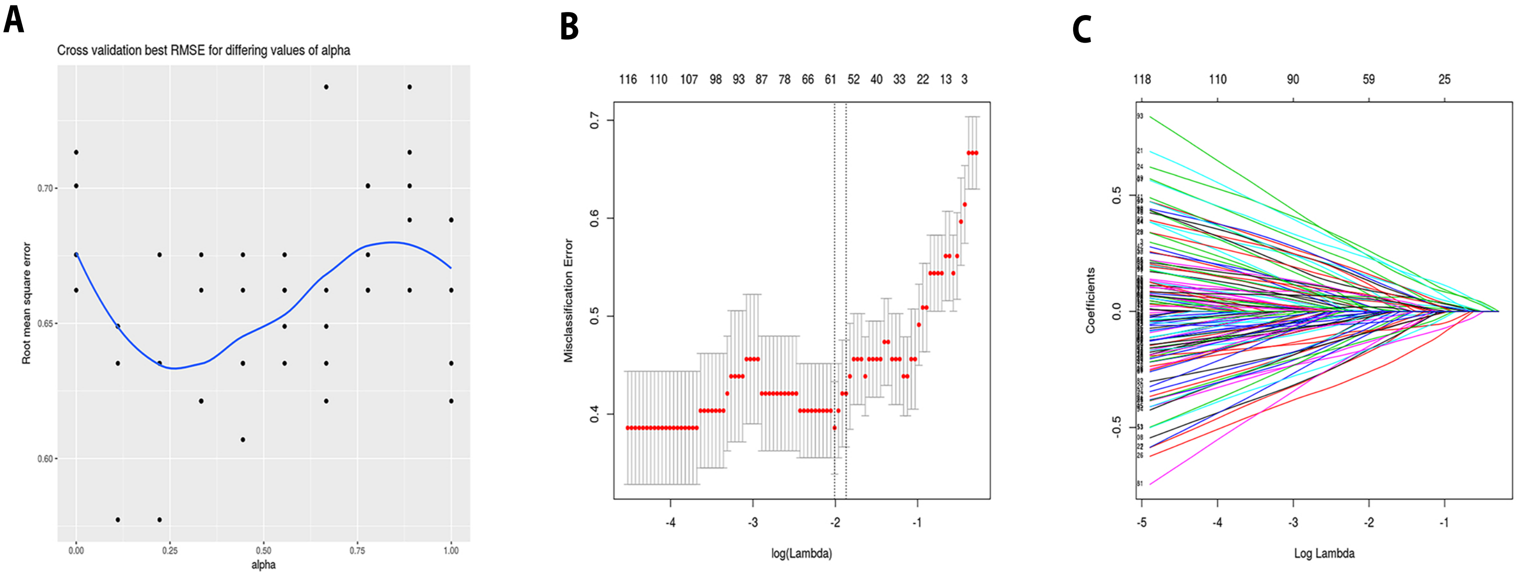


Supplementary Figure 2: Selection of metabolites using elastic net regularization. (A) Tuning alpha parameter, the parameter representing the degree of mixing between lasso (alpha=1) and the ridge regularization(alpha =0). Y-axis is the root mean square error of the 10-fold cross-validation. We selected alpha =0.22 as it gave us the minimum error. (B) Tuning lambda, the parameter controlling the shrunk rate of coefficients in the linear model. Y-axis is the misclassification error of the 10-fold cross validation. X-axis is the range of lambda, with the optimal lambda=0.008 as it gave us the minimum misclassification error. (C) The shrinkage coefficients of the metabolites using tuned alpha and lambda. Only metabolites with non-zero coefficient were be selected.
