## Supplementary material for "Pre-pregnant obesity of mothers in a multi-ethnic cohort is associated with cord blood metabolomic changes in offspring": Figure S2

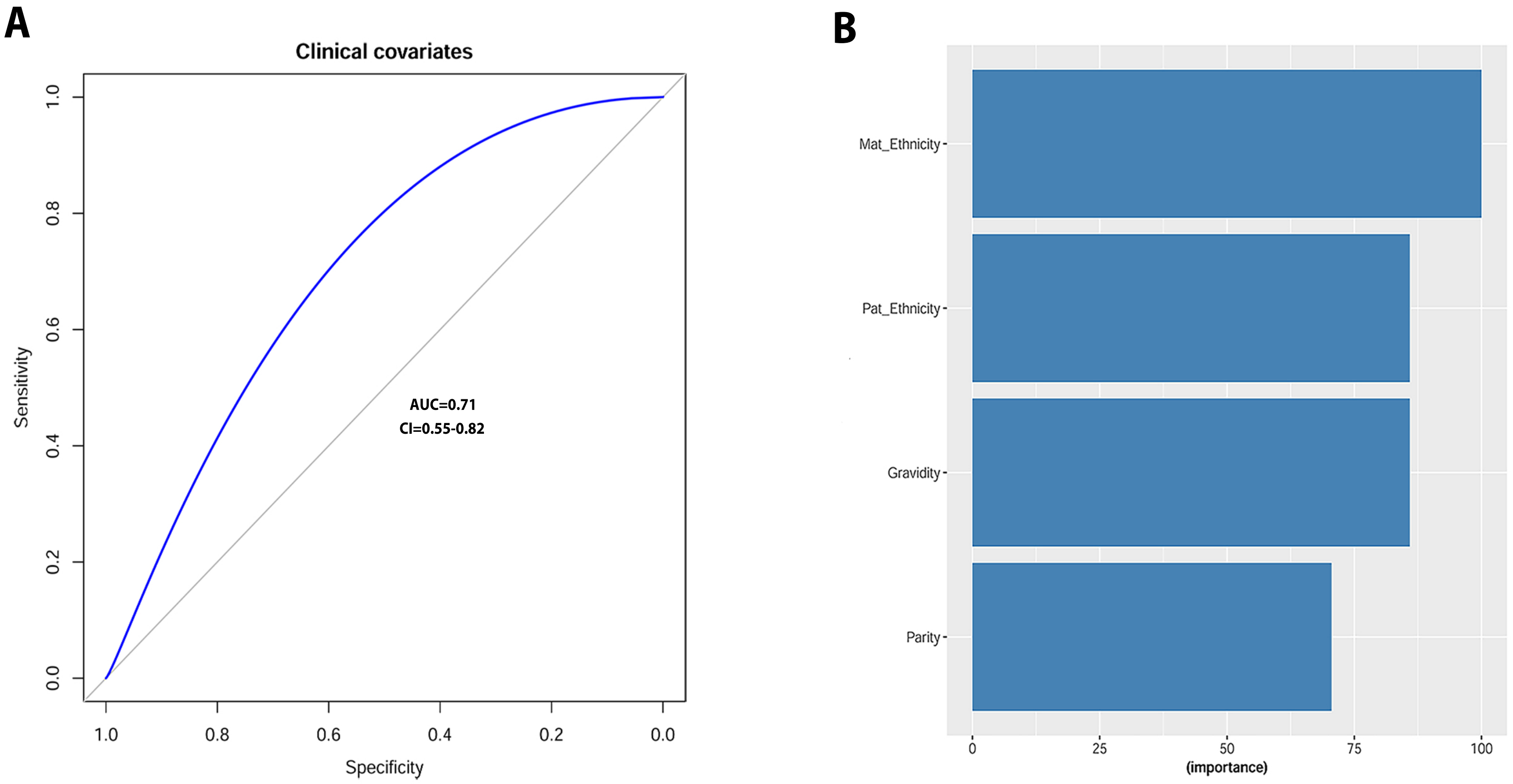


Supplementary Figure 3: Accuracies of logistic regression models and important features selected by the clinical model. (A) Model accuracy represented by classification Receiver Operator Curves (ROCs). (B) The ranking of contributions (percentage) of selected clinical features in the model.
