## Supplementary material for "Pre-pregnant obesity of mothers in a multi-ethnic cohort is associated with cord blood metabolomic changes in offspring": Figure S4

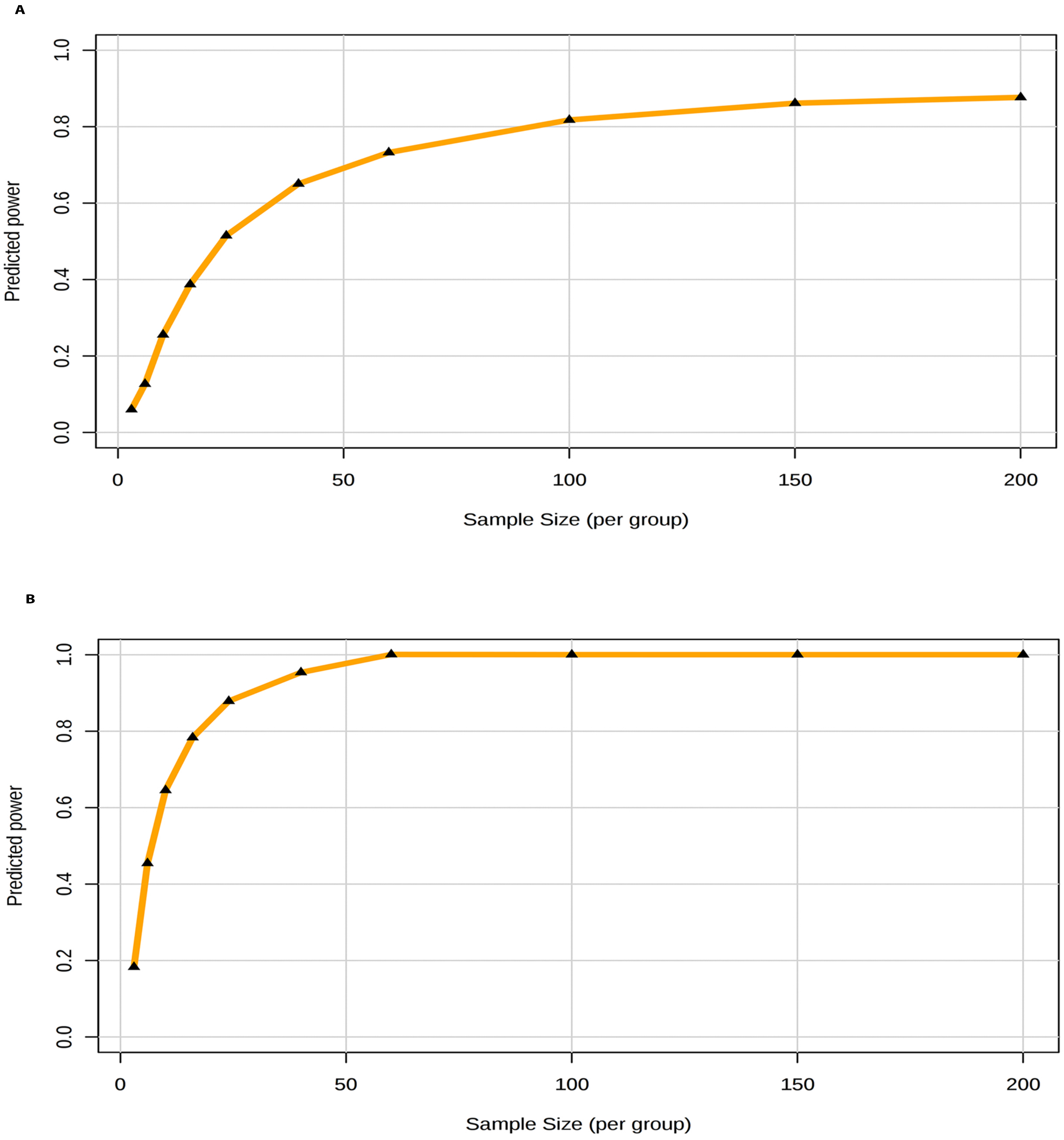


Supplementary Figure 4: Power analysis and sample size estimation plot using (A) 230 metabolites and (B) 29 metabolites that were selected by elastic-net model.
